# Metabolically-driven flows enable exponential growth in macroscopic multicellular yeast

**DOI:** 10.1101/2024.06.19.599734

**Authors:** Nishant Narayanasamy, Emma Bingham, Tanner Fadero, G. Ozan Bozdag, William C Ratcliff, Peter Yunker, Shashi Thutupalli

**Affiliations:** Simons Centre for the Study of Living Machines, National Centre for Biological Sciences (TIFR), Bangalore, India; International Centre for Theoretical Sciences (TIFR), Bangalore, India; School of Physics, Georgia Institute of Technology, Atlanta, GA, USA; Interdisciplinary Graduate Program in Quantitative Biosciences, Georgia Institute of Technology, Atlanta, GA, USA; School of Biological Sciences, Georgia Institute of Technology, Atlanta, GA, USA; Woods Hole Marine Biological Laboratory

**Keywords:** experimental evolution, multicellularity, biophysical scaffolding

## Abstract

The ecological and evolutionary success of multicellular lineages is due in no small part to their increased size relative to unicellular ancestors. However, large size also poses biophysical challenges, especially regarding the transport of nutrients to all cells; these constraints are typically overcome through multicellular innovations (e.g., a circulatory system). Here we show that an emergent biophysical mechanism — spontaneous fluid flows arising from metabolically-generated density gradients — can alleviate constraints on nutrient transport, enabling exponential growth in nascent multicellular clusters of yeast lacking any multicellular adaptations for nutrient transport or fluid flow. Surprisingly, beyond a threshold size, the metabolic activity of experimentally-evolved snowflake yeast clusters drives large-scale fluid flows that transport nutrients throughout the cluster at speeds comparable to those generated by the cilia of extant multicellular organisms. These flows support exponential growth at macroscopic sizes that theory predicts should be diffusion limited. This work demonstrates how simple physical mechanisms can act as a ‘biophysical scaffold’ to support the evolution of multicellularity by opening up phenotypic possibilities prior to genetically-encoded innovations. More broadly, our findings highlight how cooption of conserved physical processes is a crucial but underappreciated facet of evolutionary innovation across scales.

---

The evolution of multicellularity transformed life on Earth, evolving repeatedly across the tree of life ^1–3^. Size plays a central role in the early evolution of multicellularity, underpinning diverse benefits that favor a multicellular life history. For example, larger size can enable organisms to escape predation by filter feeders, increase resource utilization efficiency, and improve motility ^4^. Within more complex lineages of multicellular eukaryotes (i.e., plants, animals, fungi, and macroalgae), whose success over the last billion years has radically transformed Earth’s ecology, size plays a fundamental role in their life histories, both facilitating extensive ecological niche partitioning and underpinning the evolution of cellular and tissue-level differentiation ^2,5–8^.

The evolution of large multicellular size poses a number of constraints, however, many of which are biophysical in nature. One of the most significant of these challenges is transporting nutrients into the multicellular group. Beyond a critical size, diffusion alone is unable to transport enough resources to meet the demands of an entire group of cells ^6,9–11^. As a result, growth is typically confined to the group’s surface and, in general, the biomass increase is not exponential ^12,13^. For instance, it is well-known that microorganismal colonies exhibit sub-exponential growth beyond a certain size due to nutrient or oxygen limitations ^14–18^. Further, the absence of nutrients can affect morphology, as seen in bacterial colonies grown in various environments ^19–22^. These biophysical constraints may be mitigated by the evolution of novel biological mechanisms, such as cilia that generate fluid flows to enhance nutrient transport ^9,23^, or vascular networks that enable active transport of resources throughout the body ^5,11,24–26^. The evolution of novel multicellular transport mechanisms can lead to an evolutionary feedback loop, where the evolution of larger organism size creates steeper diffusive nutrient gradients, which favors the evolution of increasingly sophisticated transport mechanisms, thereby allowing body size to further increase. This positive feedback is thought to play a fundamental role in the evolution of large, complex multicellular organisms like plants and animals ^11^. While both theoretical predictions ^6,9–11,14,15,27^ and experiments with extant organisms ^10,28,29^ suggest that diffusion limitation is an unavoidable constraint on early multicellularity, it is difficult to address this question directly, as the early ancestors of extant multicellular lineages have long been extinct.

Using long-term experimental evolution of novel multicellularity in the snowflake yeast model system, enabling exponential growth even at macroscopic sizes beyond diffusive transport limits. To do so, we study the growth of snowflake yeast, a model system of early, undifferentiated multicellularity. Snowflake yeast have been undergoing selection for large size in the multicellularity long term evolution experiment (MuLTEE) for over 1000 daily rounds of selection ^4,17,30–33^. Within the first 600 days, they evolve macroscopic group size, where individual clusters are millimeters in diameter, and contain hundreds of thousands of clonally-related cells ^30^. We show that, beyond a threshold cluster size, the metabolic activity of snowflake yeast causes spontaneous buoyancy driven flows through the cluster that actively transport nutrients, sustaining exponential growth well beyond prior theoretical predictions. This work demonstrates how simple physical processes can act as biophysical scaffolds, opening up new frontiers of phenotypic evolution in nascent multicellular organisms even prior to the evolution of genetically-encoded innovations.

## Results

Snowflake yeast clusters grow due to the proliferation of their constituent *S. cerevisiae* cells, which undergo incomplete cell division. This causes daughter cells to remain attached to parents, thereby forming cellular chains within the clusters. Snowflake yeast remain mechanically stable even at macroscopic sizes due to the entanglement ^31^ of the cellular chains within the groups (Fig. 1**A**). These large clusters far exceed the size (≈ 50 *µ*m) at which diffusive transport alone is predicted to be sufficient to meet the cellular growth demands ^18,22,34–36^ (Supplementary Information). Further, yeast cells do not have flagella or cilia, and snowflake yeast have no known multicellular adaptations that would allow the active transport of nutrients (Fig. 1**A**, inset). The growth of these large clusters was therefore expected to be limited by the diffusive transport of nutrients deep into highly entangled clusters and thus sub-exponential. However, we found that in non-agitated nutrient liquid media, macroscopic snowflake yeast exhibits much faster and competitive overgrowth in contrast to growth on solid agar substrates (Fig. 1**B** and Supplementary Information). Indeed, we find that the growth of the macroscopic clusters immersed in a fluid environment remains exponential, even at millimetric sizes, while the growth on solid substrates becomes sub-exponential (linear) (Fig. 1**C** and Supplementary Information).

**Figure 1.**
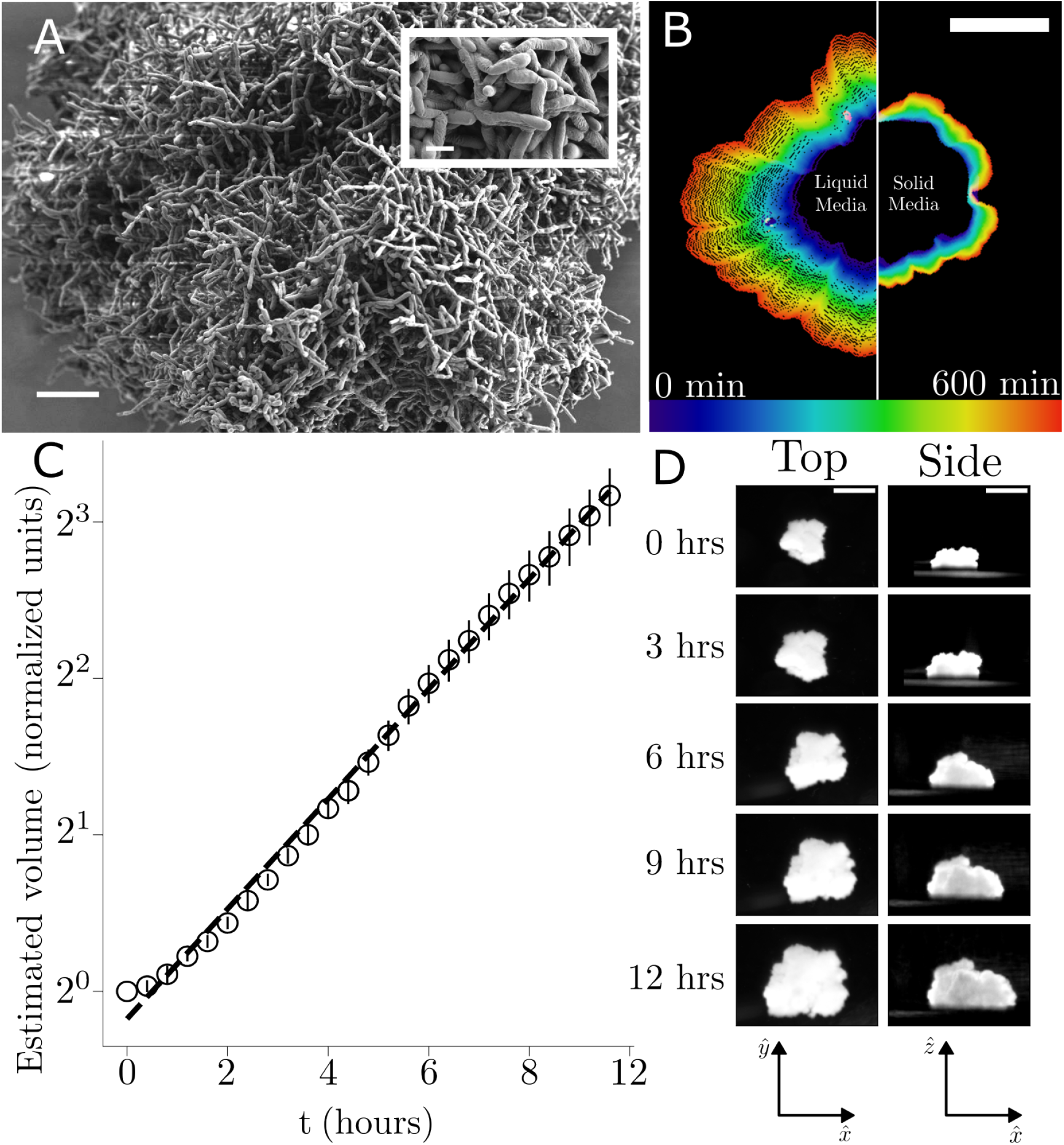
Fluid environments allow for exponential growth of the snowflake yeast clusters. **A** Scanning electron microscope image of snowflake yeast cluster (Scale bar: 20 *µ*m). *Inset:* Higher resolution image of single cells within the cluster (Scale bar: 5 *µ*m). **B** Cluster outline visualized as a function of time in non-deformable (YEPD agar) medium and fluid (YEPD liquid) environment. **C** Estimated volume (area of top view times average height of sideview) of snowflake yeast clusters over time. **D** Microscopy images of the top and side view over time of one of the clusters measured for panel C. (Scale bars: 1 mm)

These observations led us to ask how liquid media could support exponential growth of the macroscopic snowflake yeast. Given that the limits of diffusive transport were bypassed, we hypothesised the presence of an advective rather than diffusive fluid environment which could transport nutrients deep into the cluster. We found strong three-dimensional flows around the snowflake yeast (Fig. 2**A, B**). These flows had a stereotypic circulatory structure: fluid enters from the sides of the cluster, and exits from the top. Remarkably, the fastest flows around the snowflake yeast are comparable to flow speeds generated by other similarly-sized multicellular ciliated and flagellated organisms such as *Volvox* ^37,38^, choanoflagellates ^39 40,41^, *Chlamydomonas* ^37,42^, and colonial stentors ^43^ (Fig. 2**C**). These flows persist with relatively constant speeds throughout the growth of the snowflake yeast cluster (Fig. 2**D**).

**Figure 2.**
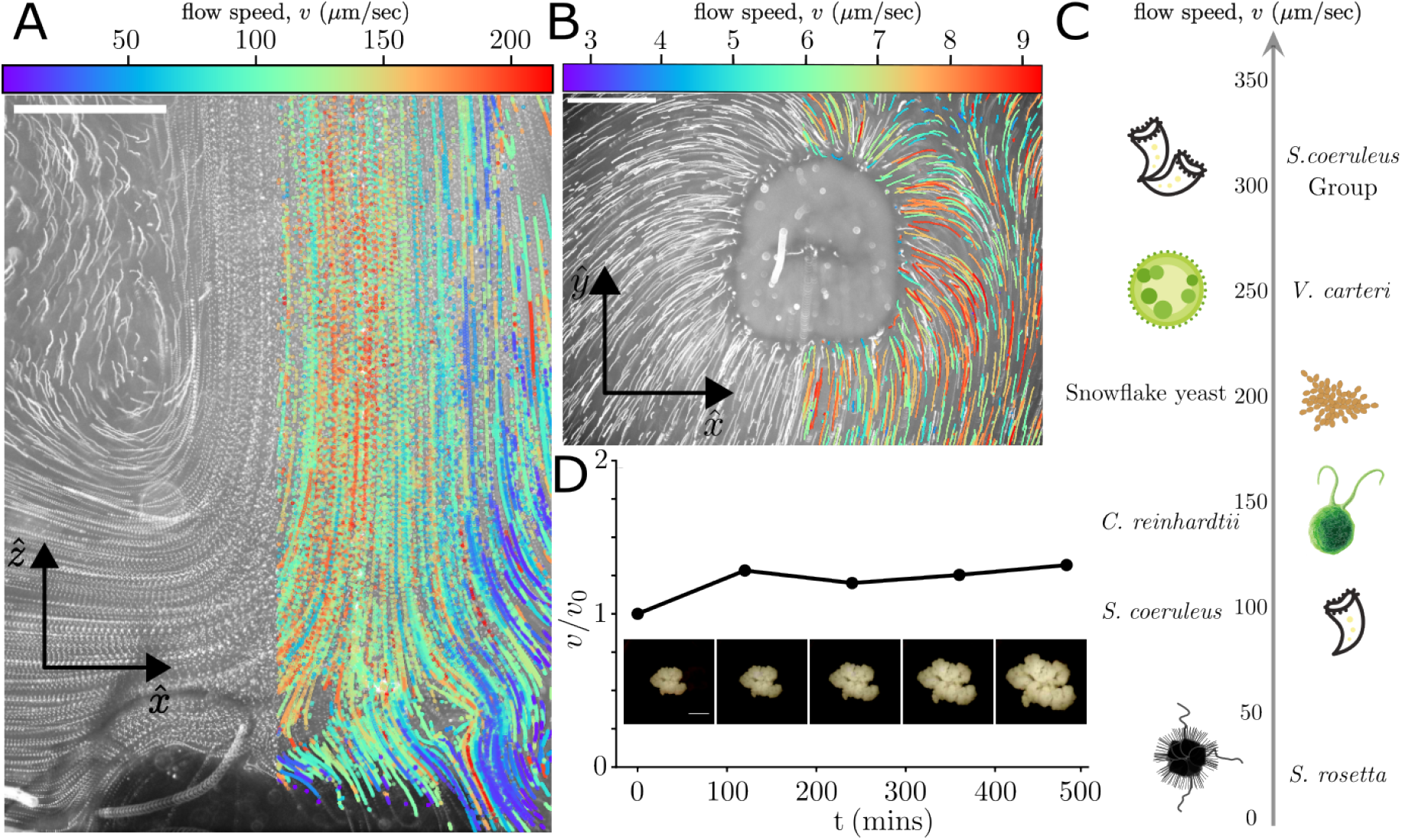
Macroscopic snowflake yeast advectively mixes its ambient fluid environment. Macroscopic snowflake yeast generate three-dimensional flows in the ambient fluid as visualised by tracer particle streaks and particle tracking data from a **A** side view and a **B** top view. The scale bar is 500 *µm*. **C** The flow speeds are comparable to those generated by ciliated and flagellated multicellular organisms. *S. coeruleus, V. carteri* and *C. reinhardtii* graphics are obtained from the Database Center for Life Sciences, CC-BY-SA 4.0. **D** The flow remains constant over a period of nearly 500 minutes during which the growth measurements were made.

While investigating the mechanism behind the flows, three observations stood out as potentially important. First, as described above, flows have a stereotyped orientation — fluid moved in at the cluster sides and upwards from the top of the cluster, and are centered on the cluster. Second, the snowflake yeast clusters we studied rely on fermentation alone for their metabolism, so in addition to depleting the surrounding media of glucose, they also produce ethanol and CO_2_, all of which are less dense than the glucose-rich media. This observation suggests that the flows are due to the generation of mass density gradients in the fluid. Indeed, previous work has shown that microbial colonies can generate flows in their surrounding fluid due to such density gradients ^44,45^. We ruled out other possibilities such as Marangoni flows, driven by surface tension gradients ^46^, as well as evaporation-driven fluid currents ^47^ (see SI for more details). Based on these observations, we hypothesized that the flows we observed were due to spontaneously generated fluid mass density gradients driven by the metabolic activity of the clusters.

To test this hypothesis, we first note that if flow is driven by a mass density gradient, we would expect the flow orientation to be sensitive to the direction of gravity. We therefore engineered a setup to measure the flow around the cluster in the same plane (the mid-plane of the cluster) in two opposing orientations of the setup with respect to gravity. In other words, we start with the cluster on the bottom of the chamber, and then flip the chamber upside down so the cluster is on the top of the chamber. In such a scenario, the direction of bouyancy driven flows in the fixed plane should reverse when the orientation of the cluster is changed. This is precisely what we observed in upon measuring the direction of the flow fields (Fig. 3**A**), thus establishing that the observed flow is sensitive to gravity and driven by the density gradient in the surrounding fluid.

**Figure 3.**
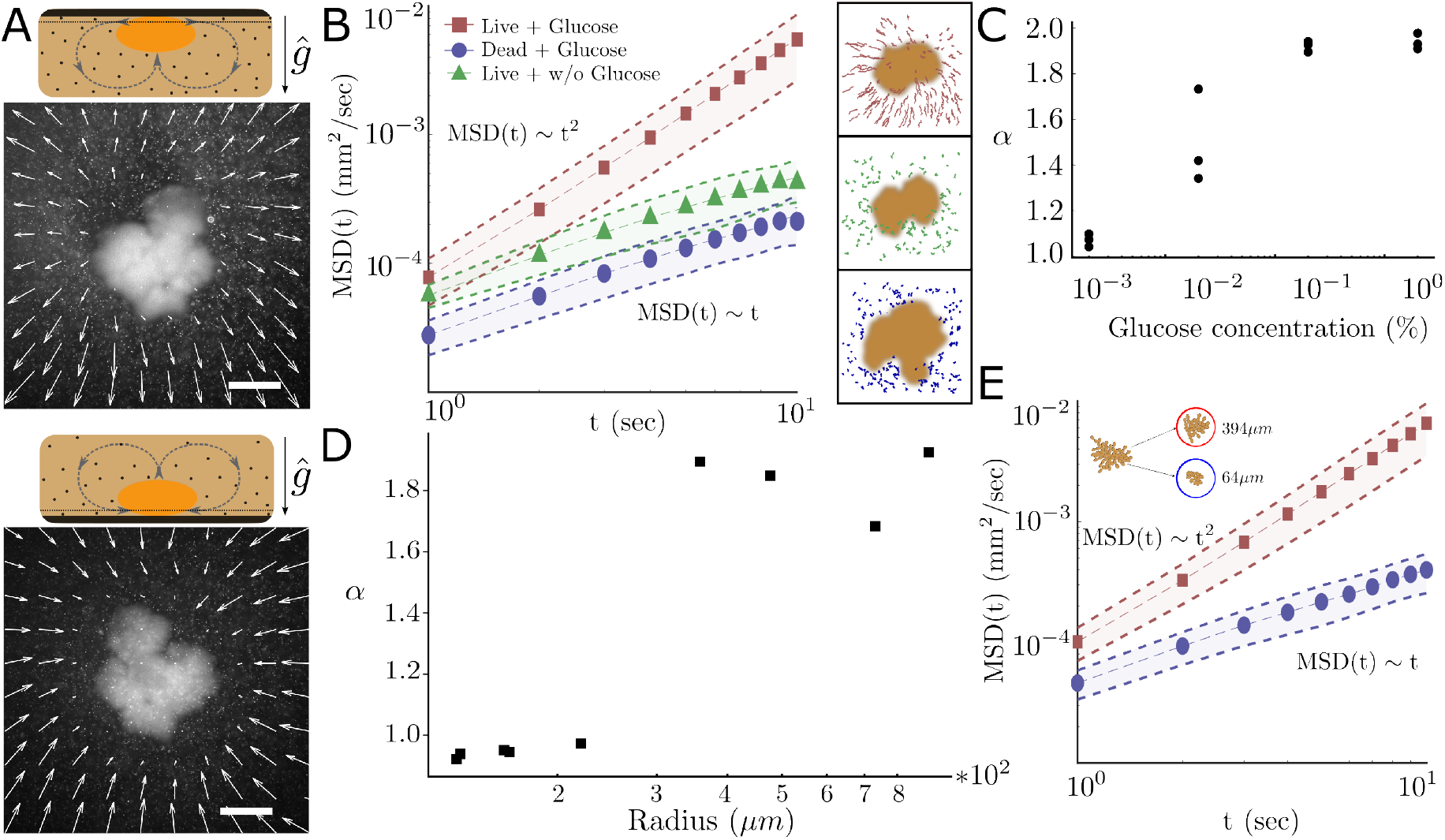
Snowflake yeast beyond a threshold size drive buoyant flows due to their metabolic activity. **A** Flows around snowflake yeast immobilised in agar (black region). The dotted line shows the imaging plane in which the flows are measured. The top and bottom panels show the flows in the same imaging plane when the experimental set-up is flipped with respect to the direction of gravity. **B** The reversal of flow in the same imaging plane hint at a gravity sensitive flow mechanism such as buoyant flows. Metabolic activity is necessary for the flows which are quantified by the mean-squared-displacement (MSD) of tracer beads around the snowflake yeast. Tracer beads around live, metabolically active snowflake yeast exhibit ballistic motion *i*.*e. α* ≈ 2 (square data points), while tracer beads diffuse, *i*.*e. α* ≈ 1, around metabolically inactive and dead clusters (triangles, circles respectively). **C** The metabolically active flows emerge at high enough glucose concentration in the ambient medium. **D** The flows emerge around clusters beyond a certain threshold size along the evolutionary lineage of the MuLTEE. The exponent *α* of the tracer particle MDSs exhibits a clear transition from a diffusive behaviour to a ballistic behaviour. **E** When clusters large enough to produce flows are broken to sizes below the threshold size identified in **D**, the clusters no longer generate flows (*α* ∼ 1, circles). On the other hand, clusters below the threshold size when aggregated together, create a flow (*α* ∼ 2, squares).

How are such fluid density gradients generated? As noted earlier, snowflake yeast metabolism relies on the uptake of nutrients (mostly glucose) from the surrounding fluid, with ethanol being produced as an outcome of fermentation. We thus placed clusters in media with and without the primary carbon source for metabolism (i.e., PBS media with or without glucose), and examined dead clusters in the presence of glucose (see SI for more details), and checked for the presence of the flows. As a quantitative test for the presence of an advective flow, we tracked the motion of micron-sized tracer particles around snowflake yeast in different media conditions. In the absence of active fermentation (i.e., due to the yeast being dead, or alive but in media lacking glucose; Methods, Fig. 3**B**), the tracer particles exhibited diffusive Brownian motion, quantified by a linear relationship in the evolution of their mean-squared-displacement (MSD) with time *i*.*e*. MSD(t) ∼ t (Fig. 3**B** and Methods). In contrast, around live clusters suspended in growth media, the MSD of the tracer particles exhibited a super-linear scaling with time; specifically, MSD(t) ∼ t^2^, indicating that they were advected because of the presence of flows (Fig. 3**B**). Further, we found that there is a threshold glucose concentration below which no flows were observed. This phenomenon was quantified by measuring MSD as a function of time and fitting to MSD ∝ t^*α*^, and extracting the best-fit power law exponent *α* (Fig. 3**C**). A value of *α* ∼ 2 is reflective of an advective environment and the tracer motion is ballistic; on the other hand, when *α* ∼ 1, tracer motion is diffusive. Based on these observations, we conclude that the metabolism of the snowflake yeast clusters drives a density-dependent mechanism to generate a circulatory flow in the ambient fluid.

If metabolism is sufficient to drive advective flows in macroscopic snowflake yeast, then why is it that we do not see such flows around all metabolically active organisms, regardless of their size? To determine if microscopic snowflake yeast also generated flows, or whether flows emerged only beyond a threshold size, we measured isolates from different time points of the MuLTEE. We found that isolates from early time points in the evolution experiment did not induce fluid flows — the tracer particles around the clusters of this size exhibit purely diffusive motion (Fig. 3**D**). Only clusters from later in the evolution experiment (*>* 300 transfers) were able to generate advective flows (Fig. 3**D**). By plotting the measured values of *α* against cluster size, we observe that *α* increases from ∼ 1 to ∼ 2 as a function of cluster size (Fig. 3**D** and Methods). This observation suggests that the metabolic activity of a sufficiently large cluster is necessary to generate these flows. To confirm if this is indeed the case, we fragmented macroscopic clusters (radii ≈ 1 mm), into smaller pieces. We found that while the smaller fragments were incapable of generating flows, fragments larger than the threshold size retained the capacity to generate flows (Fig. 3**E**). Altogether, metabolic activity mediated by sufficient nutrients in the ambient medium, together with a threshold cluster size, drives a spontaneous fluid density gradient that results in advective flows around the clusters.

Finally, while we have demonstrated that the observed advective flows arise due to metabolically driven buoyancy gradients, it is unclear if the low density fluid exhibits an instability outside of clusters. In other words, we asked whether clusters placed nearby each other would produce a single combined flow near the midpoint between them, or if individual organisms each generate their own flows. We found that clusters, even those placed only a few cluster lengths apart, produced flow towards themselves (Fig. 4**A**). This effect was quantified using particle imaging velocimetry to measure the time-averaged flow fields between two large clusters.

**Figure 4.**
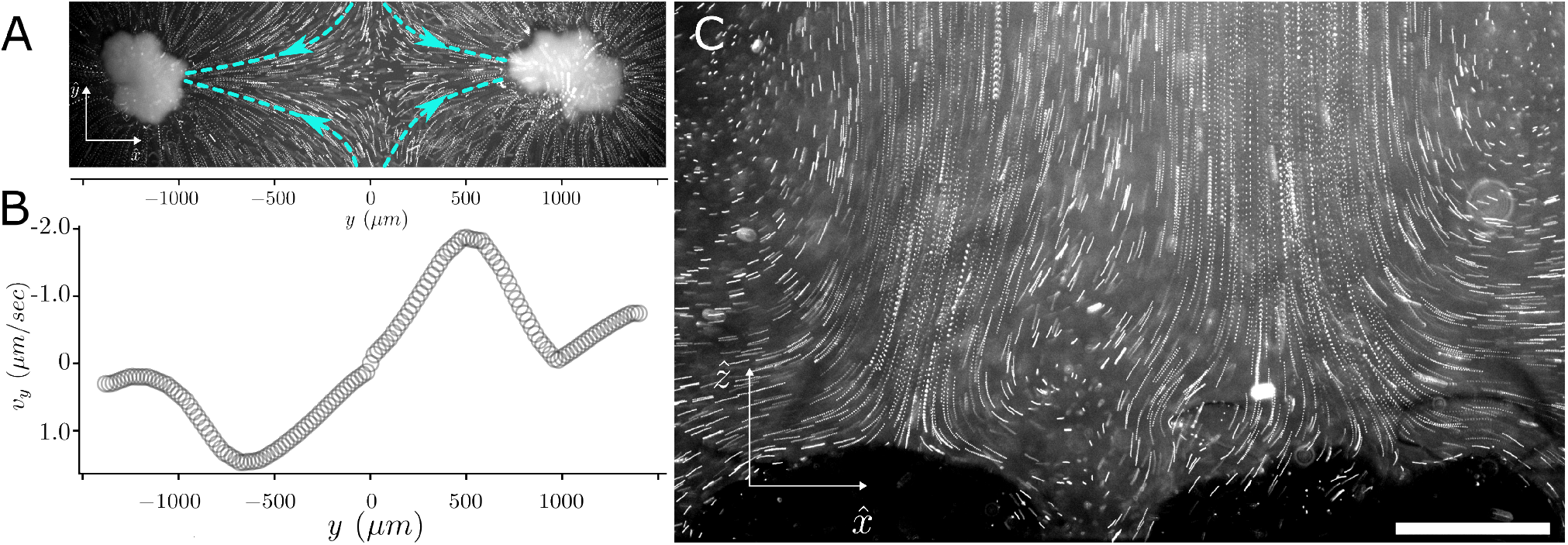
Snowflake yeast clusters act as individual metabolically powered density pumps. **A** The flows generated by the snowflake yeast are localised around individual clusters. The flows around two neighbouring clusters exhibit a stagnation point between the clusters. **B** The velocity along the line connecting the two clusters shows a singular stagnation point of the flow. **C** The presence of such a stagnation point (a PIV analysis is shown in the Supplementary Information) is also seen from the traces of the vertical plumes of two clusters that are within a cluster-radius distance from each other.

The streamlines of the flow indicate a stagnation point between the clusters. This stagnation point and flow reversal on either side of the stagnation point can be seen from the quantification of the flow velocity along a line connecting the centers of two neighbouring clusters (Fig. 4**B**). We found that the velocity dropped to zero roughly at the midpoint between the two clusters; further, the flow velocity pointed away from the center towards the clusters. This feature is seen even when the clusters are within a cluster-distance of each other (Fig. 4**C**). Altogether the flows are an emergent feature of single clusters and therefore each individual cluster acts as an independent metabolically powered density pump.

## Discussion

Growth in three dimensions poses an inherent physical challenge for multicellular organisms: as they grow in size, their surface area to volume ratio decreases, which proportionally limits the surface available for the diffusive exchange of nutrients and waste products between the organism and its environment ^11,34^. Consequently, cells located in the interior of these large organisms may be deprived of nutrients, and therefore growth limited, due to the insufficient diffusive transport of resources ^48^. This limitation is thought to impose an upper bound on the size that multicellular organisms can achieve in the absence of evolved transport mechanisms, such as vascular systems, which efficiently distribute nutrients throughout the organism ^11^. In this paper, we demonstrated that large, undifferentiated multicellular organisms can overcome diffusion limitations through emergent metabolically-driven flows. The flow emerges when the cells within clusters above a threshold size metabolize sufficiently quickly, creating a sustained density gradient in the surrounding liquid environment. Snowflake yeast are thus able to grow exponentially to macroscopic size, indicating that a consistent (i.e., size-independent) proportion of the cells are reproducing as the cluster grows.

The flows in our experiments are a consequence of yeast metabolism. There are precedents of metabolically-created density driven flow: for example, single-celled (as opposed to snowflake) yeast colonies grown in a viscous medium stir their ambient media via a spontaneously driven baroclinic instability that is due to cellular metabolism ^44^. The flows in these experiments cause the yeast colonies to break up, as the unicellular yeast are not attached to one another, ultimately disrupting the flow. Convective flows have also been seen in growing bacterial populations ^45^, which later work demonstrated is due to evaporatively-driven flows ^47^ or surface-tension driven (Marangoni) flows ^49,50^. On the other hand, bacterial biofilms have been shown to transport nutrients via structures that act like channels ^51,52^. While channels that pass through biofilms create more interfaces for diffusive flux^52^, a more complex, fractal architecture of capillaries, such as that found in the circulatory systems of animals, is needed to ensure that nutrients can actually reach every cell ^53^. Despite such transport, biofilms have not been shown to exhibit exponential growth. Additionally, biofilms typically have a dense extracellular matrix that also may prevent sufficient fluid transport through the interstices, especially where there are no channels. What we demonstrate in this paper is a much more facile mechanism that does not need to fulfill specific mathematical rules, and is independent of topologically-complex flow channels. During the MuLTEE, snowflake yeast evolved to form mechanically-durable clusters (as strong and tough as wood) via cellular entanglement ^30,31^. In addition to mechanical stability, the cellular entanglement ensures that the snowflake yeast are porous and therefore benefit, from the flows they generate, to grow up to macroscopic sizes. Altogether, the structure and porous architecture of snowflake yeast, which is a consequence of the evolution during MuLTEE, may thus play an important role in exponential growth via spontaneous density-driven flows.

This work highlights the critical interplay between physical and biological processes in the evolution of multicellularity. The spontaneous emergence of flows in snowflake yeast clusters demonstrates how a purely physical mechanism, arising from the basic physical laws governing fluid dynamics, can profoundly impact the development of a biological system. These flows act as a “biophysical scaffold”, enabling a key trait — the ability to overcome diffusion limits — without requiring any dedicated structural adaptations. In the snowflake yeast model system, biophysical mechanisms have previously been shown to underpin key steps in the transition to multicellularity, such as the origin of a life cycle via packing-induced strain ^54^ and the emergence of heritable multicellular traits via a growth pattern guided by maximum entropy ^55^. Here, we extend these results to show that behaviors once thought to require sophisticated adaptations may instead arise ‘for free’, as a result of the emergent biophysics of simple multicellular systems. This suggests that the inherent physical properties of biological systems may have been crucial in enabling the evolution of novel multicellular traits, by allowing access to novel phenotypes that can subsequently be refined and stabilized by selection. Indeed, it will be interesting to examine how fluid dynamics affects the subsequent evolution of multicellularity in the snowflake yeast model system. Behaviors affecting flow may become genetically assimilated ^56,57^, and yeast evolved in static media may evolve novel multicellular morphologies that increase flow-based nutrient transport.

The evolution of multicellularity has long been thought to be constrained by fundamental physical limitations, chief among them being the diffusive transport of nutrients, which becomes increasingly inadequate at larger organismal sizes ^9,55^. Strikingly, we found that an entangled morphology, which first evolved to provide mechanical stability to snowflake clusters growing in rapidly shaking media ^31^, has the serendipitous side effect of enabling snowflake yeast to grow large enough to spontaneously generate circulatory flows via a widespread biophysical mechanism — buoyant instabilities triggered by localized metabolism. This work demonstrates how tradeoff breaking innovations can arise through the cooption of conserved biophysical mechanisms, with latent physical processes opening up new frontiers of phenotypic evolution when harnessed by newly-evolved traits. This observation fits into an emerging view that the interplay between physical and biological processes is both common and highly impactful in shaping evolutionary trajectories ^58,59^, and highlights the critical role of biophysical interactions in the origin of new levels of biological organization.

## Methods

### Yeast strains

We used evolved isolates of snowflake yeast from anaerobic line 5 of the Multicellular Long-Term Evolution Experiment (MuLTEE) ^30^. For almost all experiments (exponential growth, flow fields, metabolism assays), we used the 1,000-day evolved strain from line 5 (PA5 t1000). For the cluster size experiments in Fig. 3, we used 200-, 400-, 800-, and 1,000-day evolved strains from line 5 (PA5).

### Cell culture

The yeast were cultured in Yeast Extract Peptone Dextrose (YEPD) media (1% yeast extract, 2% peptone, and 2% dextrose) at 30^°^C in a shaking incubator at 250 RPM.

### Quantifying cluster area during growth

A single cluster was placed in liquid YEPD (see cell culture methods) in one well of a 12-well plate. A 45-degree mirror (Thor labs right-Angle Prism Dielectric Mirror, 400-750 nm, L = 10.0 mm) was placed in the chamber next to the cluster, making the side profile of the cluster visible. Each of these clusters was imaged every 30 minutes for 12 hours on a Zeiss AxioZoom.V16 microscope. Both the top of the cluster and the side of the cluster (via the mirror) were imaged at every timepoint. The clusters were kept at room temperature during this time. For analysis, clusters were segmented using the connected components algorithm in the scikit-image library with Python 3.10. The total number of pixels of the segmented clusters was found.

### Measuring flow fields

To measure flow fields, we filled a shallow well (WPI’s FluoroDish tissue culture dishes, with a well 1.2 mm tall and 23 mm in diameter) with YEPD (or other liquids, depending on conditions being tested) and placed one large snowflake yeast cluster in the well. The well was then covered with a cover slip larger than the well diameter to seal it. Images were taken at a rate of one image every 5 seconds on a Zeiss Axio Zoom.V16 microscope or Olympus XI81 inverted microscope. Presence and absence of advective tracks were quantified using particle tracking with the Mosaic plugin on the image analysis software ImageJ. To measure plumes in the XZ plane, we used a 45-degree angle mirror (MRA12-E02, ThorLabs) placed in one well of a 12-well plate with 3 ml of YEPD media and a single (or multiple) clusters. Images were taken at a rate of at least 1 fps. The flow velocities in Fig. 2 were quantified using the TrackMate software in Fiji/ImageJ with an LoG spot detector and a Kalman filter for tracking. The tracks were thresholded for quality, track length, and maximum speed to remove spurious tracks.

### Literature review for flow speeds

To generate Fig. 2**C**, we searched the literature for flow fields of ciliated or flagellated microorganisms and small multicellular organisms. We took the maximum flow speed, if provided in the text of the papers, or if not, we took the highest recorded speed from charts of the flow fields. For snowflake yeast, we took an estimate on the high end of the vertical flow speeds as quantified by particle tracking (described above) in Fig. 2**A**.

### Gravitational flow field assay

To measure the effect of the axis of gravity on the orientation of the flow, we made a chambers using PDMS a flexible silicone polymer. These chambers were circular in shape with a diameter of ≈ 800 *µ*m and height of ≈ 500 *µ*m. In order to adhere the cluster to the one side of the chamber we used a mixture of 0.5% agar in phosphate buffered saline. Just before the agar became stiff, we placed a single cluster of ≈ 300 *µ*m radius atop the agar. Once the cluster was fully adhered (approximately 1-2 minutes), we filled the chamber with YEPD nutrient media supplemented with 0.5 *µ*m RFP-coated polystyrene beads. We placed a cover slip atop the chamber and sealed it with nail polish. This allowed us to invert the entire chamber without dislodging the cluster.

### Protocol for breaking clusters

In order to break the clusters without damaging them, we used a cut pipette tip and pipetted a single cluster from the t1000 population (≈ 1000 *µ*m). Via the mechanical action of the pipetting, we were able to break the single cluster into smaller clusters of ranging from ≈ 50 *µ*m to ≈ 600 *µ*m radius. We allowed the resulting liquid culture to settle under gravity for 40 seconds. This resulted in most of the larger clusters settling to the bottom of the tube. By pipetting from either the surface or the bottom of the liquid, we were able to recover small clusters of ≈ 50 *µ*m radius and large clusters of ⪆ 300 *µ*m radius.

## Acknowledgments

We acknowledge Yash Rana and Aditya Iyer for initial observations of the flow. Gabriela Canales, Yuan Wang and Celia Cui helped explore these dynamics during the 2022 and 2023 Physiology courses at the University of Chicago’s Marine Biology Lab (Woods Hole, MA). We thank the Department of Atomic Energy (India), under project no. RTI4006, and the Simons Foundation (Grant No. 287975), the Central Imaging and Flow Cytometry Facility, and the computational facilities at the NCBS. We also acknowledge support from the National Institutes of Health (U.S.A.) under grant numbers NIH T32GM142616, R35GM138354 and R35GM138030.

## Supplementary Information

**Figure S1.**
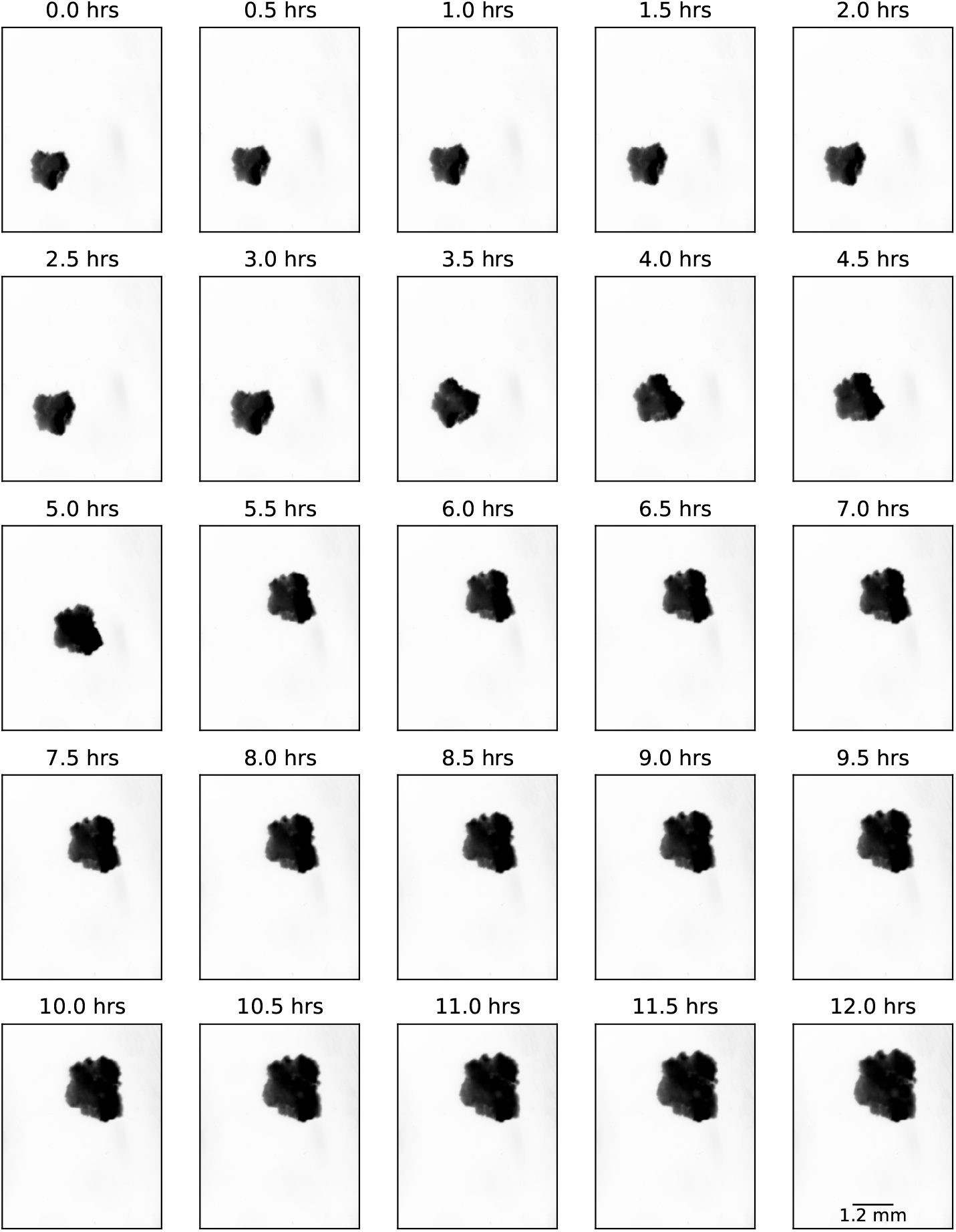
Growth of a snowflake yeast cluster over 12 hours. Grown in liquid YEPD. Viewed from the top.

The information in this document is arranged according to the figures presented in the main text of the paper. This includes detailed protocols, data acquisition and analysis methods.

### Figure 1

#### Panel A

Snowflake yeast clusters from PA5 t1000 were grown in YEPD medium till stationary phase, harvested and washed in double distilled water three times. The clusters were next diluted in increasing concentrations of ethanol till final concentrations of 100 percent ethanol was reached. Clusters were super-critically dried in a Leica EM CPD300 Critical Point Dryer. Once dried, the clusters were sputter coated with gold at 20m amps for 90 seconds to achieve a gold coating of ≈5 nm. Finally clusters were imaged on a Zeiss Merlin compact electron microscope. No processing was done to the resultant images.

**Figure S2.**
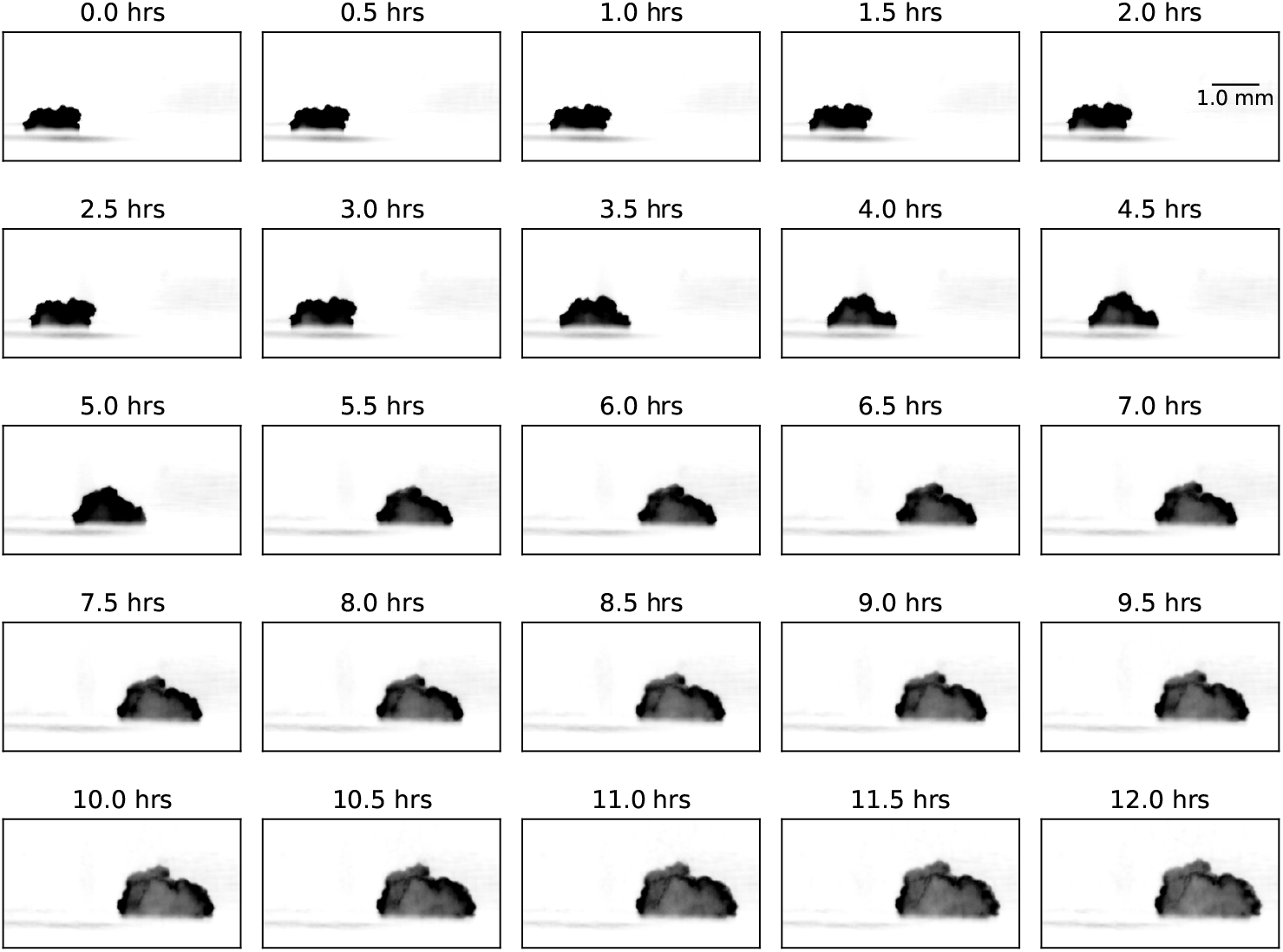
Growth of a snowflake yeast cluster over 12 hours. Grown in liquid YEPD. Viewed from the side with a 45-degree angle mirror.

**Figure S3.**
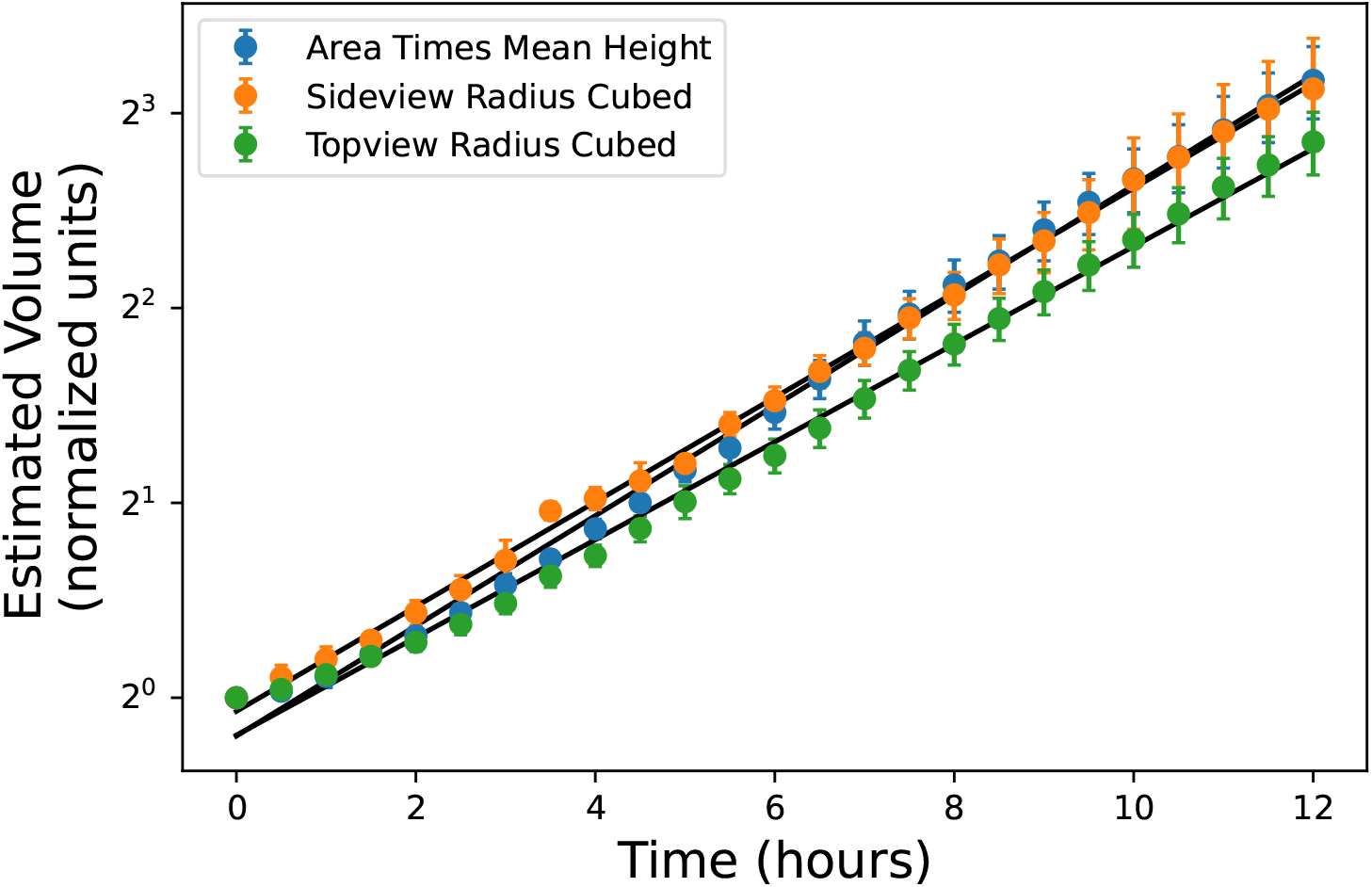
Three different volume estimates for the growth of snowflake yeast clusters.

#### Panel B and Fig. 1 S6

Two different media conditions were tested — YEPD in 2% agar and YEPD in water. In order to prepare YEPD agar plates, yeast extract (1%), peptone (2%) and agar (2%) were dissolved in 475 ml distilled deionized water and sterilized by autoclaving. Following this, 25 ml of 40% filter sterilized glucose was added to achieve a final glucose concentration of 2%. 3ml of this medium was poured into individual 35 mm Petri dishes and left to set for 30 minutes. Once set, in order to ensure as little fluid on top of the agar as possible, the plates were further allowed to dry for 2 hours prior to usage.

Preparation of liquid YEPD medium was exactly the same as mentioned for agar without the addition of 2% agar. Before pouring 3 ml of the liquid YEPD into a petri dish, a thin layer (1 ml) of 0.5% agar in phosphate buffered saline was poured into each 35 mm Petri dish. This layer of 0.5% agar in phosphate buffered saline served as a pad to prevent the clusters from moving once inoculated, and since it does not contain any nutrition, it does not affect the interpretation of our results.

**Figure S4.**
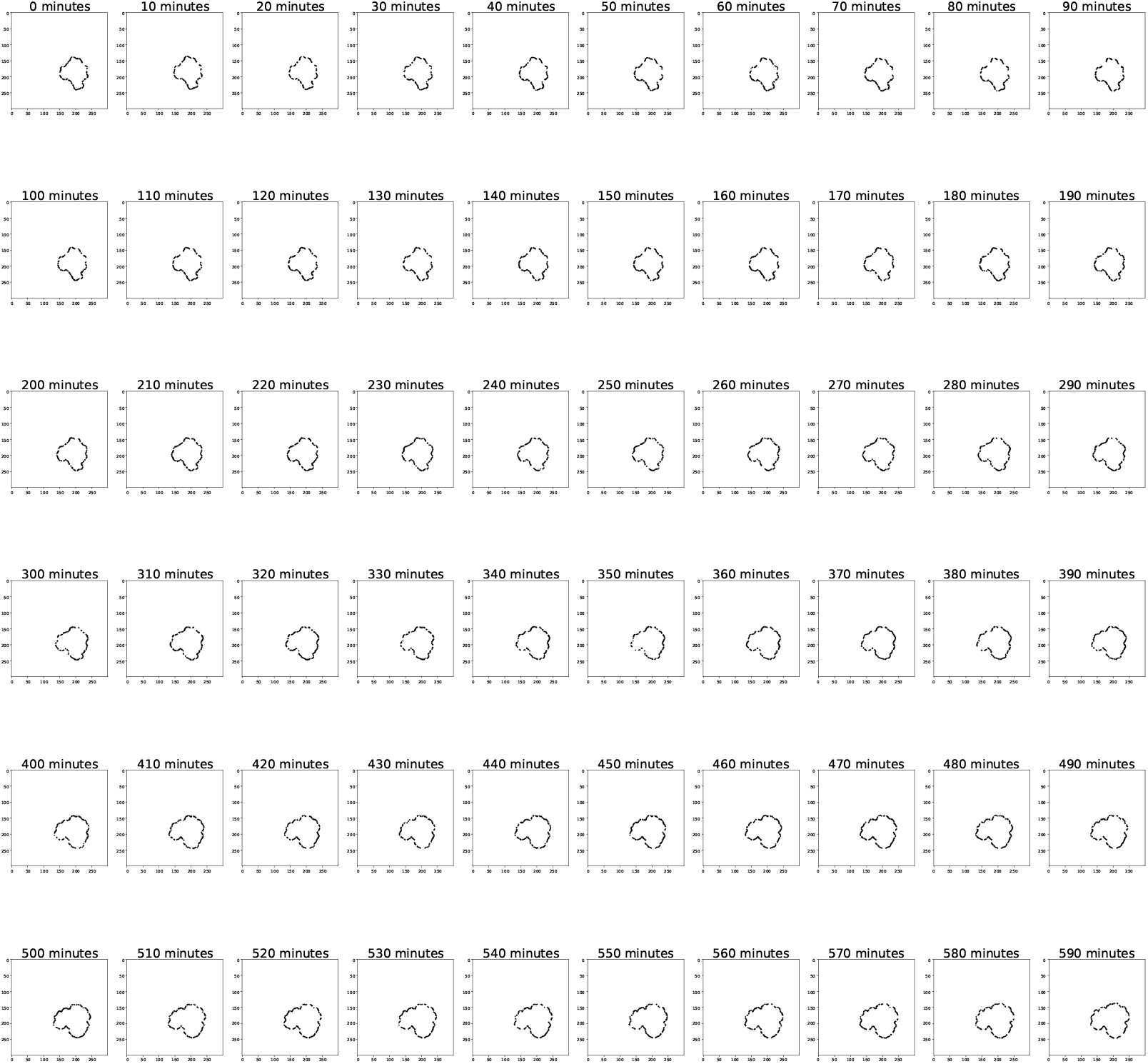
Growth of snowflake yeast over 10 hours in 2% agar.

Plates containing either liquid YEPD or YEPD agar were inoculated with single clusters which had been isolated and washed thrice in PBS following growth in YEPD for ≈24 hours. For single cell experiments, similar preparatory steps were taken with the only difference being that, following the last wash, cells were pelleted and 2 *µ*l of the pellet was used to inoculate each YEPD agar plate. (Single cell experiments could not be performed in the same way as those performed for clusters in liquid YEPD, as single cells would diffuse, preventing accurate imaging of the sample.)

Plates were next placed on an Epson v800 flatbed scanner and scanned every ten minutes for 10 hours at a resolution of 1200 dpi. The ambient temperature and relative humidity of the room in which imaging was done was 25^°^C and < 60%.

#### Panel C

PA5 t1000 clusters were grown for 24 hours in 10 ml of YEPD in a shaking incubator at 30 C and 250 RPM. Single large clusters were obtained for the experiments described in this section by taking 1 ml out of this 24 hour culture, letting it settle for around 30 seconds, and pipetting out most of the supernatant to remove smaller clusters, and pipetting in fresh YEPD. This washing was repeated several times to ensure the removal of any small clusters. Then around 500 ml of this mixture was pipetted into an empty Petri dish with a few ml of YEPD in order to spread out the large clusters into a larger surface area so that individual clusters would be visible to the eye and available to select for experiments. A chosen large cluster is carefully pipetted up into a wider-bore tip 1000 *µ*l pipette tip, and this tip is gently placed into fluid. The cluster is allowed to sink down the pipette and out into the waiting fluid (without actively pipetting it out).

**Figure S5.**
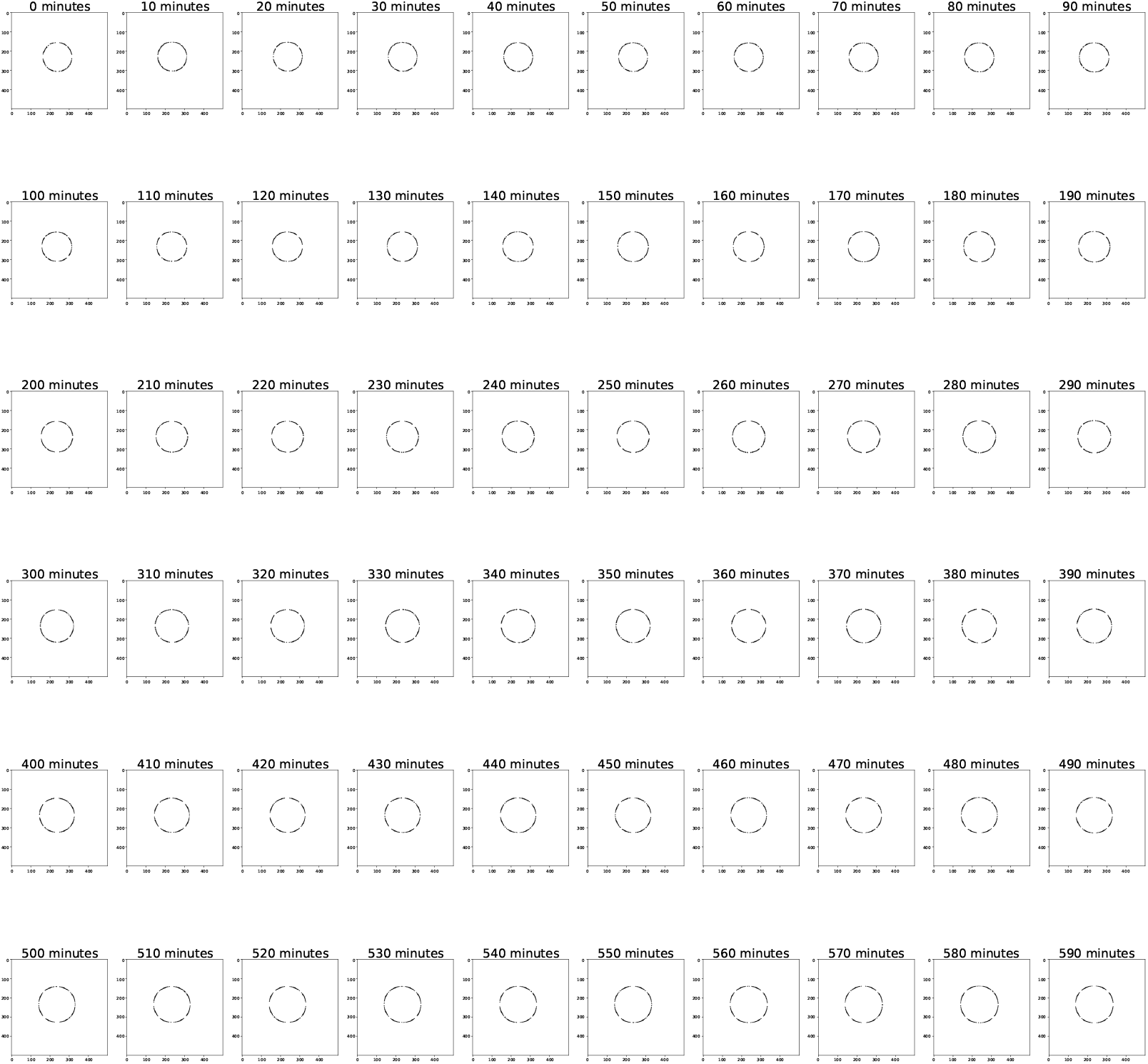
Growth of single-celled yeast over 10 hours in 2% agar.

A single PA5 t1000 yeast cluster was placed in 3 mL of liquid YEPD in one well of a 12-well plate. A 45-degree mirror (Thor labs right-angle Prism Dielectric Mirror, 400-750 nm, L = 10.0 mm) was placed a few millimeters from the cluster. Images were taken of the top of the cluster and the side of the cluster (via the mirror) every 30 minutes for 12 hours using a Zeiss AxioZoom.V16 microscope. Three replicate measurements were obtained.

The clusters were segmented by binarizing and using a connected components algorithm in the scikit-image Python package. The estimated volume was calculated by multiplying the area of the topview (calculated as the number of pixels multiplied by the area scale factor) by the average height of the sideview (calculated by finding the height in pixels of each column from top to bottom, averaging, and multiplying by the distance scale factor). SI Fig. S3 compares this estimate to two other estimates: the cube of the maximum radius of the sideview, and the cube of the maximum radius of the topview.

**Figure S6.**
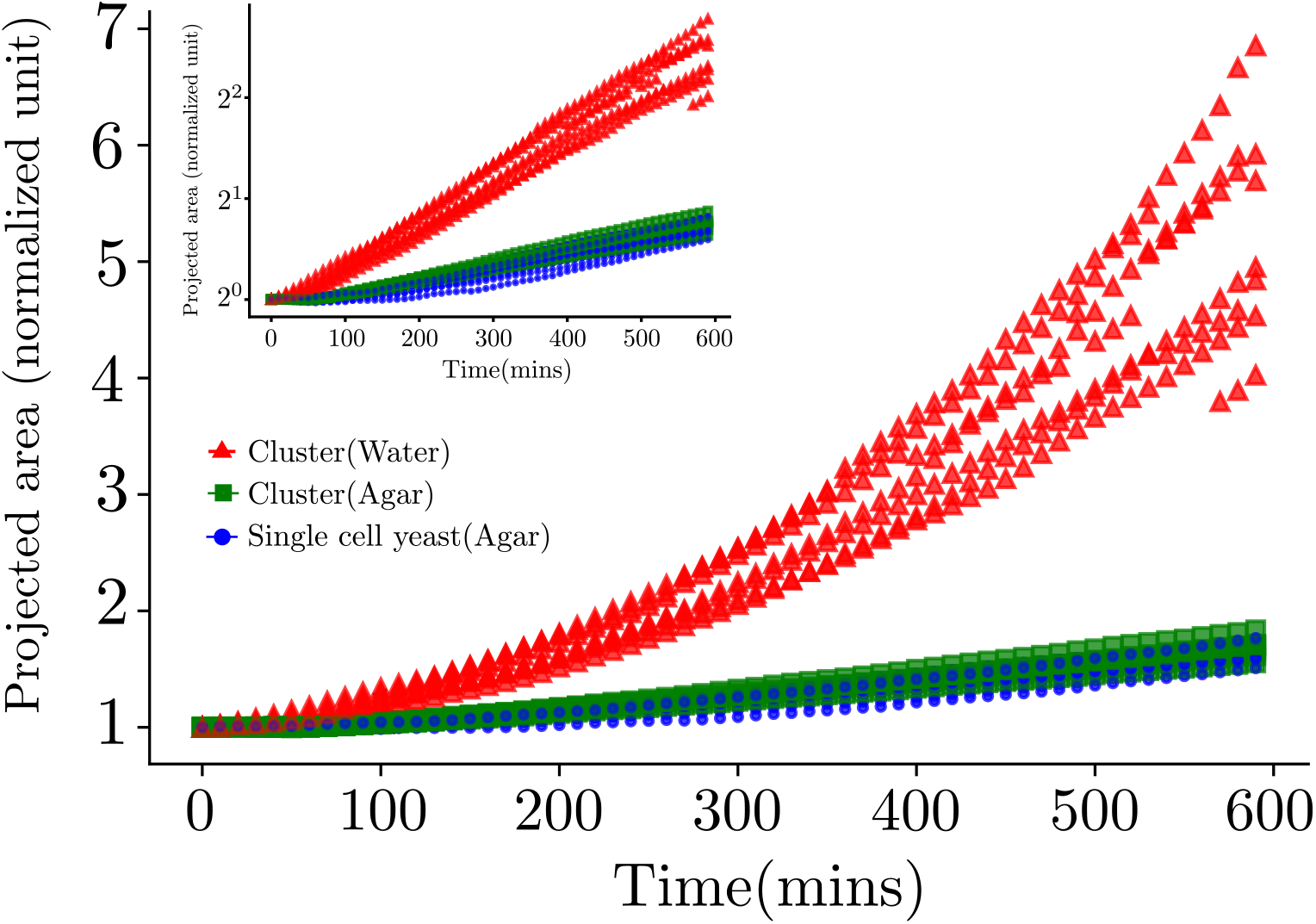
Comparison of growth of snowflake clusters in YEPD (liquid) and YEPD (agar) and single-celled clusters in agar.

### Figure 2

#### Panels A and B

PA5 t1000 clusters were grown overnight in YPED media at 30^°^C. To image a cluster, the cluster was placed in a small Petri dish with a 45-degree mirror (Thor labs right-Angle Prism Dielectric Mirror, 400-750 nm, L = 10.0 mm), and images were taken on a Zeiss AxioZoom.V16 microscope at approximately 3 fps for both the top view of the cluster and the side view in the mirror.

Particle tracking was done using the TrackMate plugin in FIJI with a LoG spot detector and a Kalman filter for connecting spots. Exact parameters can be found in the supplemental data. The tracks were filtered for spot quality, track length, and max speed to remove as many spurious tracks as possible. Tracks were overlaid and visually compared to the video to check that they were capturing the motion of the tracing particles.

A composite image of the video was made by taking the max of every fourth frame, creating an effect similar to a star tracks image in order to show the particle movement. The particle tracks were then overlaid on the right side of the image. The colorbar indicates the maximum speed of each track.

See Figs. S7 and S8 for additional examples of particle tracking of flows, and see the supplementary videos for the corresponding videos.

#### Panel C

A literature search was performed for flow fields of small organisms, especially ciliated and flagellated organisms. Most reports found included a chart displaying the flow field around the organism, from which we obtained the approximate maximum speeds of the flow fields. The value used for snowflake yeast is the approximate maximum value from Panel A. The flow speeds found around snowflake yeast clusters can vary based on the cluster, the chamber geometry, and other factors.

#### Panel D

Single snowflake yeast clusters were assayed across 10 hours for the presence or absence of ambient fluid mixing in pseudo 2D chambers. These chambers were prepared by sticking multiple layers of double sided tape, with each tape having a height of ≈0.1 mm (chamber heights were adjusted depending on the size of the clusters being observed, ranging from 0.2 mm - 1 mm) on a previously cleaned glass slide. PA5 t1000 clusters were grown for 2 hours in YEPD liquid medium, following which they were washed thrice in phosphate buffered saline and inoculated into the chamber along with YEPD liquid supplemented with 0.5 µm GFP coated beads. Presence or absence of advective mixing was assayed every hour by taking a 2-minute video at a frame capture rate of 5 fps in the GFP fluorescence channel.

**Figure S7.**
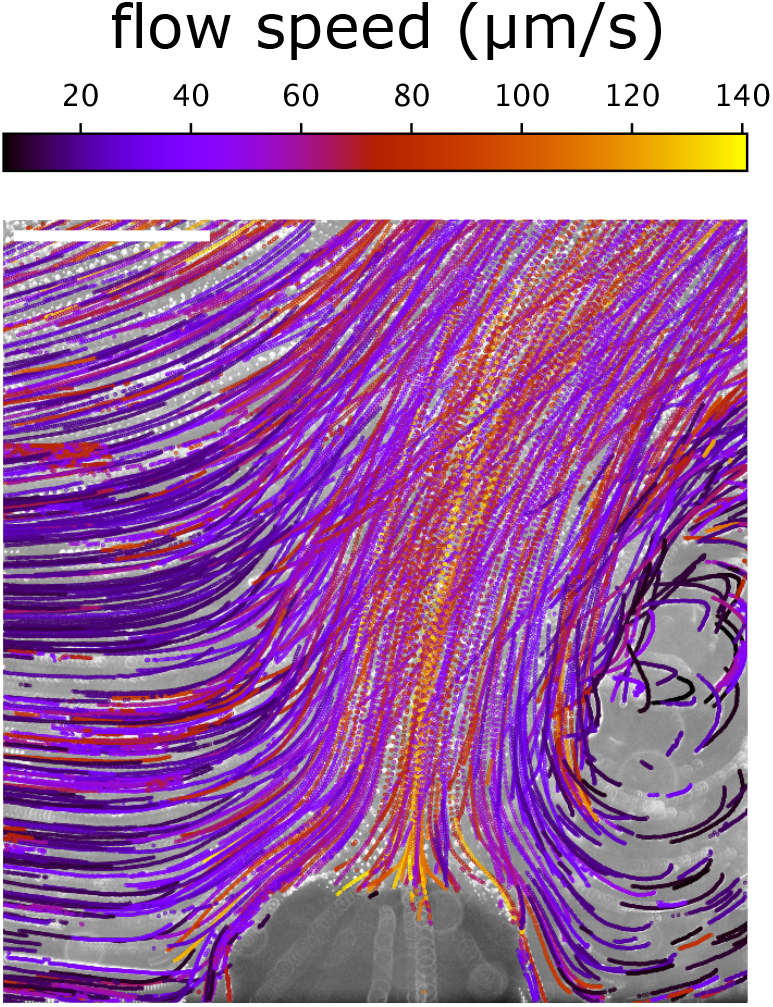
Additional example of particle tracking for the X-Z view of a cluster (viewed with a 45-degree mirror). Scale bar: 500 µm.

**Figure S8.**
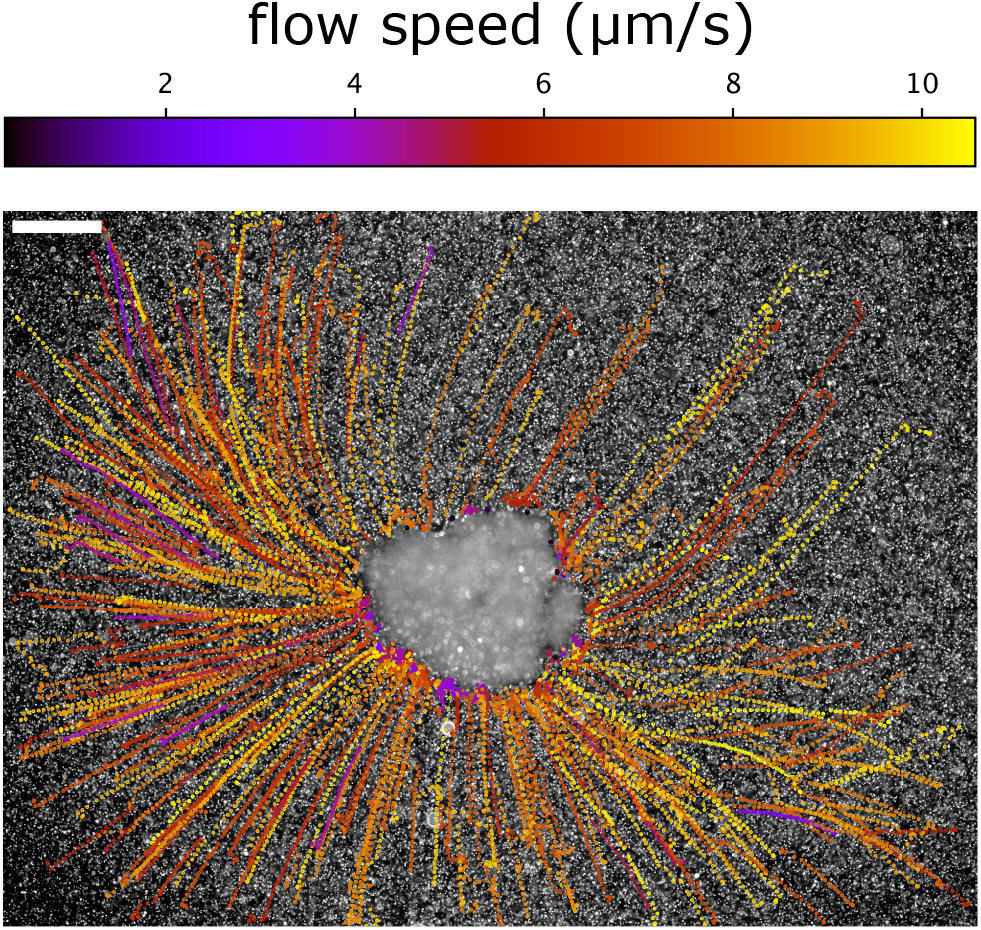
Additional example of particle tracking for the X-Y view of a cluster. Scale bar: 500 *µ*m.

The video generated was loaded into image analysis software FIJI, where, following conversion to 8-bit grayscale, particles were detected and traced using the Mosaic particle tracker plugin. Detected particles were then analyzed in Matlab to calculate the mean squared displacement (MSD) of individual particles across each time step. These MSDs were then plotted against the time step in order to quantify the presence or absence of ambient fluid mixing. A slope of ≈ 1 on a log log MSD vs. time step plot corresponds with a diffusive fluid environment and a slope of ≈ 1 corresponds with a advective fluid environment. The velocity of the mixing is the intercept of this graph and was plotted against the time points. The velocity was normalized since the magnitude of the mixing is dependent on the geometry of the chamber and varies depending on the imaging condition.

### Figure 3

#### Panel A

A chamber was engineered that could be inverted completely without any leaks, and more importantly, without clusters to dislodging from the the position in which they were inoculated. In order to make the chamber, a silicone polymer known as poly dimethyl siloxane, PDMS for short, was used. This polymer, when treated with the appropriate curing agent, hardens into a flexible and transparent rubber. Approximately 10 ml of this polymer was allowed to cure in a 60 mm Petri dish, which resulted in a circular rubber of 60 mm radius and ≈5 mm height. Once cured, individual square shaped pieces of ≈10 mm side length were cut which were then punched with a 5 mm biopsy punch in order to create a circular hole with 5 mm radius and 5 mm height. Next the cut piece of PDMS and a previously cleaned coverslip were plasma cleaned on one side. Once plasma cleaned, the piece of PDMS was bonded onto the coverslip, resulting in a circular well of 5 mm radius and 5 mm height. Into this well 200 *µ*l of 0.5% agar in PBS was added in order to adhere clusters to one side of the chamber. Within 10 seconds of the addition of the base agar, a single PA5 t1000 cluster was inoculated atop the base agar. The chamber was next filled with liquid YEPD supplemented with 2% glucose or PBS supplemented with 2% glucose (roughly 500 *µ*l) and 0.5 *µ*m GFP tracer particles. Once full, another previously cleaned 20 mm coverslip was placed on the open side of the well and secured with clear nail polish. This setup was left to dry for 10 minutes. Sealing the chamber with nail polish was done to ensure that no evaporative flows were generated. Imaging was done on this setup both with its right side up and also with the upside down. We confirmed that the cluster had not dislodged from the point at which it had been adhered by checking the change in the z-focus position. Imaging was done at both the first plane at which the cluster came into focus (bottom) and the last plane after which the cluster was out of the focal plane (top). The microscope used was a IX81 Olympus Epifluorescence microscope. Imaging was done at 4X magnification in the GFP fluorescence channel at a frame capture rate of 5 fps.

#### Panel B

Different media conditions were assayed to check for presence or absence of flows. Pseudo 2D chambers were prepared by sticking multiple layers of double sided tape, with each tape having a height of ≈0.1mm (chamber heights were adjusted depending on the size of the clusters being observed, ranging from 0.2 mm - 1 mm) on a previously cleaned glass slide. Once the chambers were prepared, Snowflake yeast clusters (anaerobic line 5 t1000) grown for 24 hours in YEPD were harvested and washed three times in PBS, following which they were transferred to a vial containing the medium being tested supplemented with 0.5 *µ*m GFP fluorescent beads (available in table). Individual clusters were taken from the vial containing the medium being tested and placed into a chamber. Once inoculated, a coverslip (20 X 20 mm) was placed on top of the chamber, this was then sealed with transparent nail polish which was allowed to dry for ≈10 minutes. Sealing the chamber with nail polish was done to ensure that no evaporative flows were generated. Imaging was done at magnifications of 10X and 20X in the GFP fluorescence channel at a frame capture rate of 5 fps.

#### A. Panel C

Presence or absence of flows was assayed as detailed above for different concentrations of glucose in phosphate buffered saline.

| Media | MSD slope | Replicates |
| --- | --- | --- |
| YEPD (live) | 1.739 | 3 |
| PBS + glucose (live) | 1.840 | 3 |
| PBS (live) | 1.004 | 3 |
| PBS + glucose (dead) | 1.003 | 3 |

#### Panel D

We used various time points from one of the experimental evolution lines (available in table) and assayed them for the presence or absence of ambient fluid mixing (as mentioned in SI for Fig. 3 panel B). We only used one media condition for this experiment (PBS supplemented with 2% glucose). Chamber heights were adjusted depending on the size of the cluster being assayed.

| Radii | MSD slope |
| --- | --- |
| 913.1073486 | 1.9762 |
| 225.895301 | 0.9736 |
| 139.3004523 | 0.9378 |
| 137.1524991 | 0.92 |
| 482.9431768 | 1.8957 |
| 737.078953 | 1.7217 |
| 364.827388 | 1.9441 |
| 169.6425654 | 0.9447 |
| 166.0494746 | 0.9504 |
| 190.9950942 | 0.9584 |

#### Panel E

An experiment was devised where PA5 t1000 clusters (>1 mm radius) were pipetted with a cut tip to generate different sized clusters. Clusters were independently assayed for the presence or absence of ambient fluid mixing (as described in SI for Fig. 3 panel B).

#### Fig 3 SI 1

Experiments were performed with the ancestral population from the anaerobic line 5 (which had on average radii smaller than the threshold radii) were grown as colonies on YEPD 2% agar plates, following which a piece of the colony greater than the threshold size was picked and assayed for the presence or absence of ambient fluid mixing (as described in SI for Fig. 3 panel B (main text)). Similar experiments were also conducted for single cell colonies.

**Figure S9.**
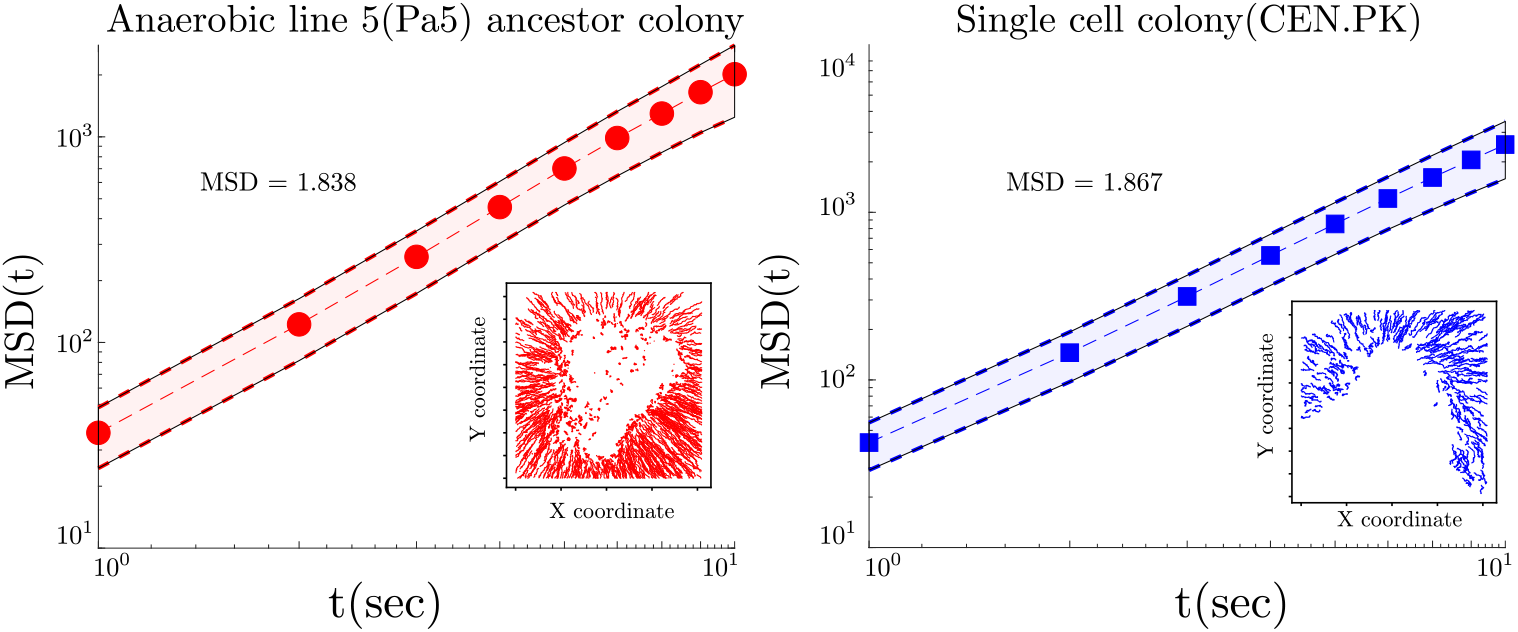
Comparison of growth of snowflake clusters in YEPD (liquid) and YEPD (agar) and single-celled clusters in agar.

### Fig 4

Multiple snowflake yeast clusters from anaerobic line 5 t1000 were observed to understand how ambient fluid mixing is affected by the presence of more than one cluster. Growing and washing steps for the clusters are exactly as mentioned in Fig. 2 panel D. Observations were made in 35 mm Petri dishes with 1 ml of 0.5% agar in PBS to adhere clusters to the point of inoculation. Following adhering, 3 ml of YEPD liquid supplemented with 2 *µ*m GFP coated particles were added. To prevent Marangoni flows due to evaporation 2 ml of squalene was added atop the fluid layer. This oil floats atop water and is biocompatible, hence it doesn’t interfere with our interpretation of the results. The setup was imaged on a Leica stereo microscope in the GFP fluorescence channel at a magnification of 1.9X with a frame capture rate of 0.1 fps.

